# The long non-coding RNA, CyKILRb, augments oncogenic phenotypes via induction of PIK3R2 and activation of the PI_3_K/AKT axis

**DOI:** 10.1101/2025.10.13.682173

**Authors:** Xiujie Xie, H. Patrick Macknight, Amy L. Lu, Charles E. Chalfant

**Affiliations:** Department of Medicine, Division of Hematology & Oncology, University of Virginia, Charlottesville, VA, 22903; Research Service, Richmond Veterans Administration Medical Center, Richmond, VA, 23249; Program in Cancer Biology, University of Virginia NCI Comprehensive Cancer Center, Charlottesville, VA, 22903

**Keywords:** Long noncoding RNAs, RNA splicing, CyKILR, PI3K, AKT, non-small cell lung cancers

## Abstract

Our laboratory recently identified a novel long noncoding RNA termed CyKILR that has two splice variants with distinct cellular localizations and opposing roles in tumorigenesis. The cytoplasmic variant, CyKILRb (exon 3 exclusion), promotes tumorigenesis, whereas the nuclear variant, CyKILRa (exon 3 inclusion), functions as a tumor suppressor. In this study, the molecular mechanism of the tumorigenic role of CyKILRb was characterized. Specifically, deep RNA sequencing analysis revealed that CyKILRb regulated the PI_3_K/AKT signaling pathway to block downstream tumor suppressors. In particular, downregulation of CyKILRb induced the loss of PIK3R2, an activator of PI_3_K, as well as RPS6KB2 and GNB2, two implicated tumor promotors, with a concomitant increase in the tumor suppressors, CDKN1A (p21) and CDKN1B (p27). In contrast, CyKILRb ectopic expression produced the opposite effect, and suppression of either PIK3R2, PI_3_K or AKT attenuated CyKILRb-induced cell proliferation and clonogenic survival. CyKILRb negatively regulated CyKILRa expression, which was blocked by inhibition of either PI_3_K or AKT. PIK3R2 ectopic expression overcame the cellular effects of CyKILRb downregulation, but not PI_3_K or AKT inhibition orienting the signaling pathway from CyKILRb→↑PIK3R2→PI3K→AKT→↓CyKILRa→enhanced oncogenicity. These findings highlight the critical role of CyKILRb in tumorigenesis and define a novel feed-forward regulatory mechanism linked to alternative RNA splicing.

## INTRODUCTION

Lung cancer remains the leading cause of cancer-related mortality worldwide, with non-small cell lung cancer (NSCLC) accounting for approximately 85% of these cases[1, 2]. Despite advancements in targeted therapy and immunotherapy, NSCLC remains a highly aggressive malignancy with limited treatment options for patients. In recent years, non-coding RNAs (ncRNAs) have emerged as critical functional regulators of gene expression, significantly influencing tumor progression and therapy resistance in human malignancies including NSCLC [3, 4]. The complexity of ncRNAs, particularly their interactions within the tumor microenvironment, adds another layer of challenge to NSCLC treatment[5]. Among ncRNAs, long non-coding RNAs (lncRNAs) have garnered increasing attention for their role in modulating oncogenic and tumor suppressor pathways through diverse cellular mechanisms including chromatin remodeling, transcriptional regulation, and post-transcriptional modulation[6, 7]. Additionally, lncRNAs exhibit complex secondary and tertiary structures, enabling them to interact with DNA, RNA, and proteins to modulate cellular pathways at multiple levels[8].

Several lncRNAs have been implicated in NSCLC progression with some functioning as oncogenes and others as tumor suppressors. For instance, MALAT1 is a well-characterized oncogenic lncRNA that promotes metastasis by regulating epithelial-mesenchymal transition (EMT) and enhancing tumor cell migration[9]. Conversely, MEG3 functions as a tumor suppressor by interacting with p53 and inhibiting cell proliferation and survival[10, 11]. LncRNAs are also involved in modulating drug resistance in NSCLC. HOTAIR, for example, contributes to resistance against tyrosine kinase inhibitors (TKIs) in EGFR-mutant NSCLC by epigenetically regulating drug-response genes[12]. Similarly, LINC00460 has been reported to promote cisplatin resistance by modulating the PI_3_K/AKT signaling pathway[13]. Given their crucial role in oncogenesis and therapy resistance, lncRNAs present promising targets for therapeutic intervention in NSCLC. However, further research is needed to elucidate the precise molecular mechanisms governing their function, structural dynamics, and regulatory interactions.

Despite their functional significance, only a small subset of lncRNAs has been identified and investigated in NSCLC as noted above, leaving many other lncRNAs uncharacterized as well as aspects of their regulatory roles unexplored[14]. In this regard, previous work from our laboratory identified CyKILR, a novel lncRNA expressed in NSCLC with wild-type CDKN2A and STK11 genes[15]. This lncRNA presented with two splice variants, CyKILRa and CyKILRb, based on inclusion/exclusion of exon 3 that exhibit opposing functions in NSCLC[15]. While CyKILRa functions as a tumor suppressor, CyKILRb promotes oncogenesis and was required to maintain tumorigenic capacity in NSCLC cells with the aforementioned oncogenotype. Still the precise molecular mechanisms underlying the roles of these lncRNA splice variants remained unclear.

In this study, we sought to elucidate the oncogenic mechanism of CyKILRb and its regulatory relationship with CyKILRa. Through deep RNA sequencing, CyKILRb was discovered to modulate a crucial signaling cascade that governs cell survival, proliferation, and apoptosis resistance, specifically the PI_3_K/AKT pathway. Furthermore, our findings revealed this novel regulatory axis controls the expression of CyKILRa via suppression of alternative RNA splicing. The feed-forward amplification mechanism identified in this study underscores the complex interplay between lncRNA splice variants in tumorigenesis and may offer future strategies for precision-based cancer treatment in NSCLC.

## RESULTS

### The cytoplasmic lncRNA, CyKILRb, induces the PI_3_K/AKT signaling pathway

Our laboratory recently reported that the lncRNA, CyKILRb, was a potent tumor promoting factor. To delve into the mechanism of action of this lncRNA, the transcriptomic signatures related to CyKILRb expression were interrogated using an unbiased approach. Specifically, CyKILRb was downregulated in two NSCLC cell lines (H1299 and H2009) followed by deep RNA sequencing and pathway analysis. KEGG pathway enrichment analysis revealed that the PI_3_K/AKT signaling pathway exhibited the most significant gene expression changes upon CyKILRb silencing (**Figure 1A-C**). Additionally, five significantly altered genes were identified based on the highest percent change and lowest p-value: downregulation of PIK3R2, RPS6KB2, and GNB2 with upregulation of CDKN1A (p21) and CDKN1B (p27). These effects were validated at the RNA and protein levels (**Figure 1D-F**) along with loss of AKT activation/phosphorylation (**Figure 1E,F**). Ectopic expression of CyKILRb in these NSCLC cell lines (**Figure 2A**), in contrast, modulated these proteins in the opposite direction as those observed with CyKILRb downregulation (**Figure 2B&2C**). These data demonstrate that CyKILRb is a specific activator of the PI_3_K/AKT axis, which is linked to modulations in PIK3R2, PRK6B2, GNB2, p21, and p27.

**Figure 1.**
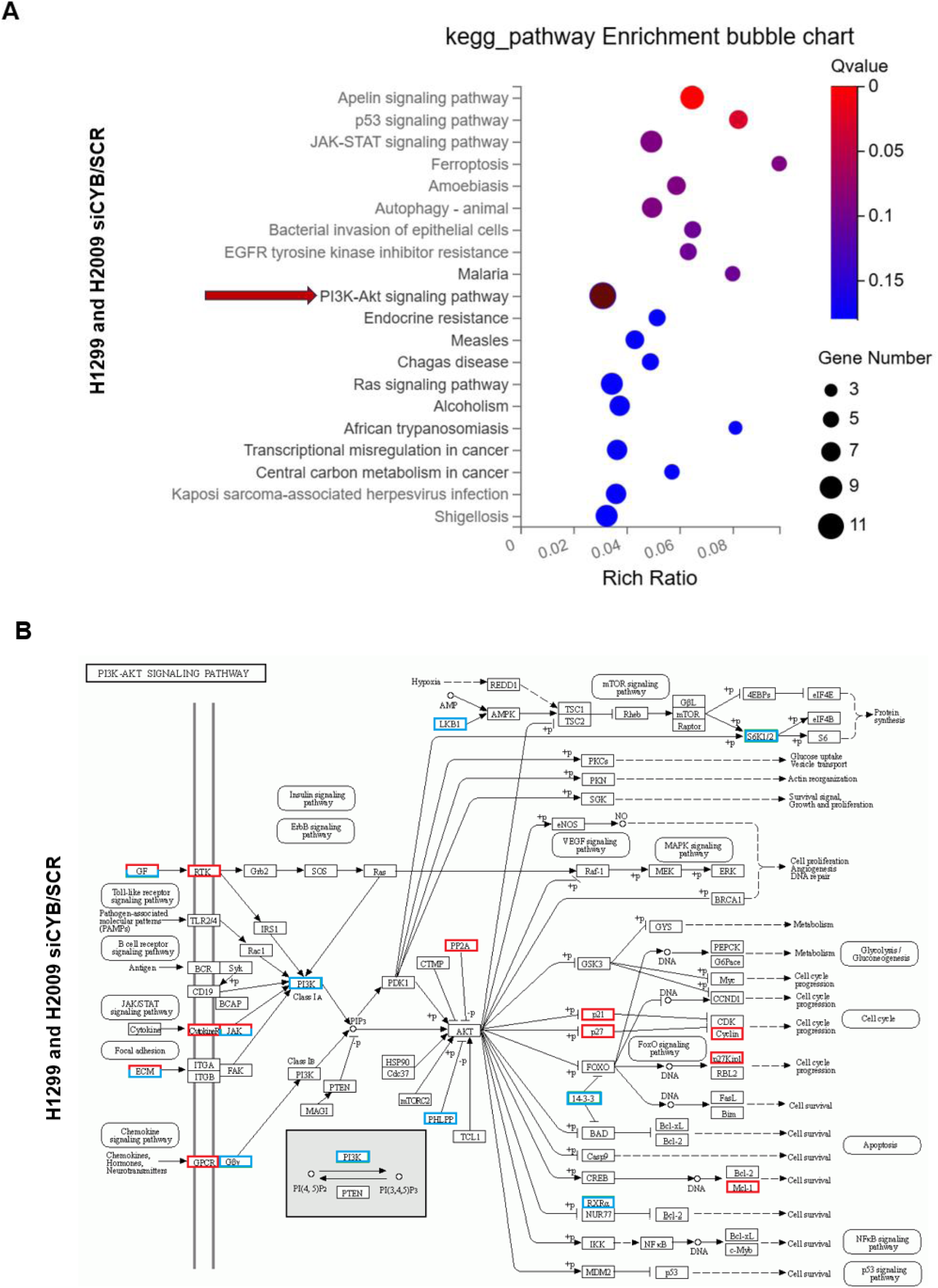
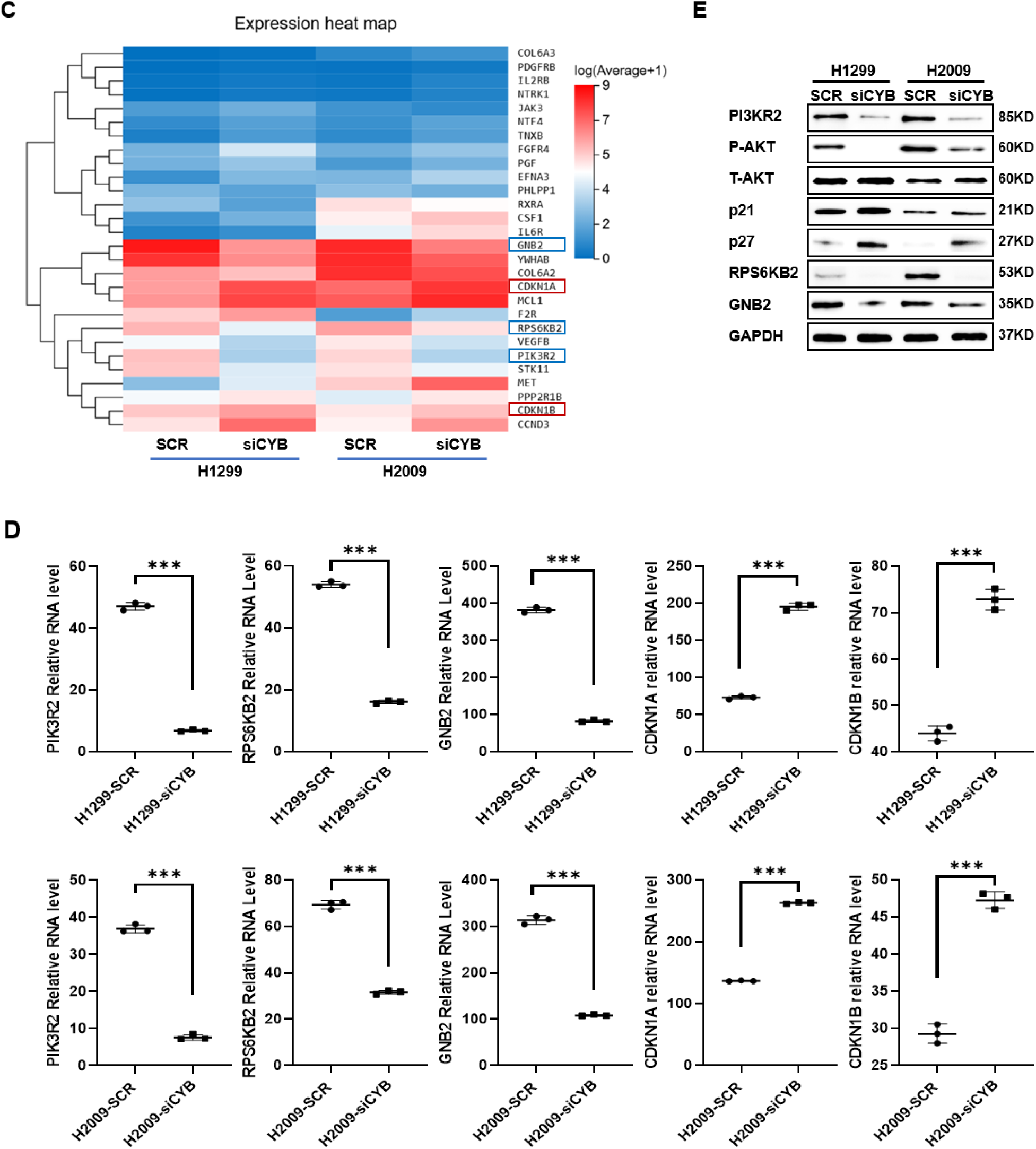
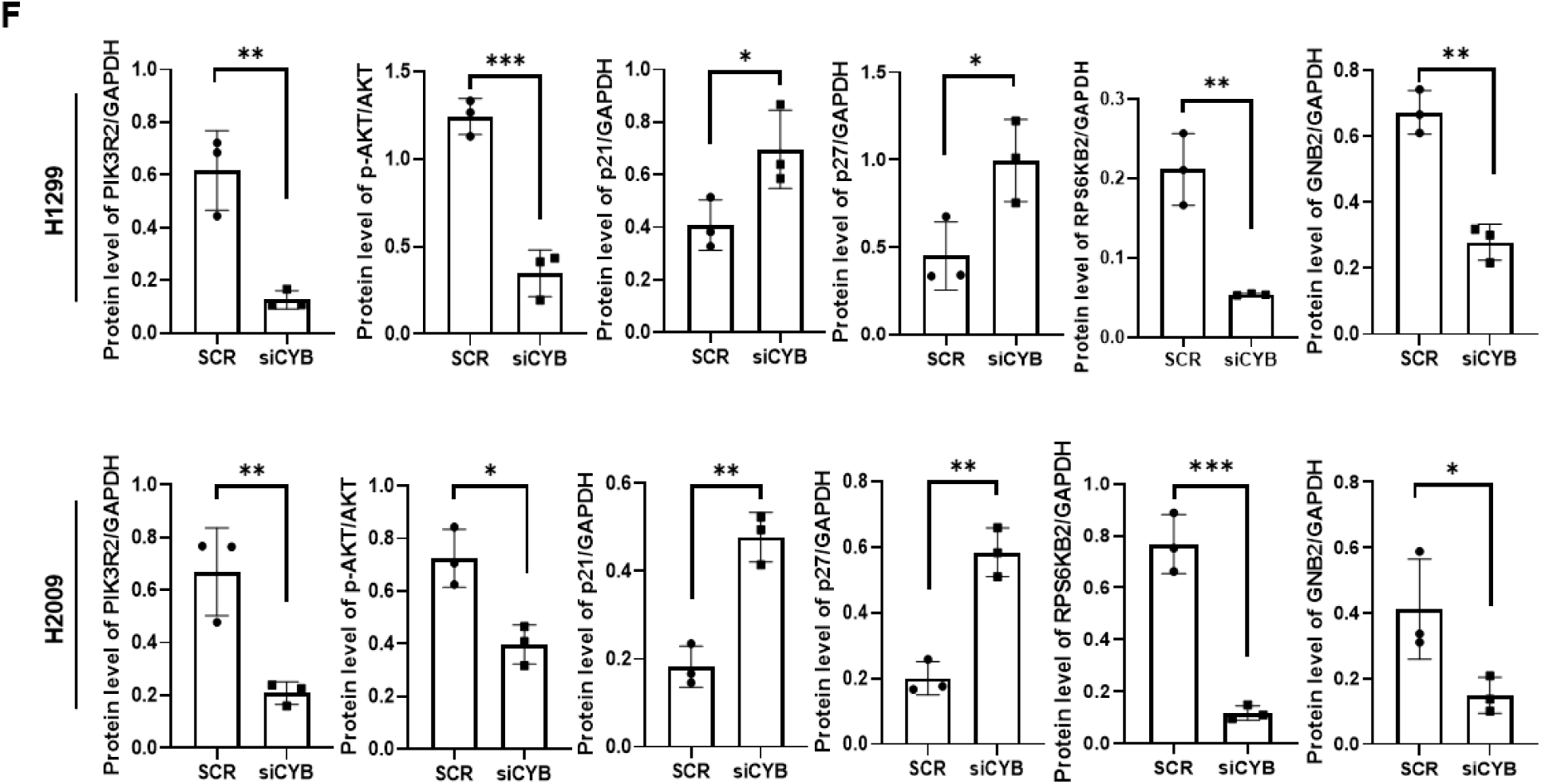
The cytoplasmic lncRNA, CyKILRb, augments the PI_3_K-AKT axis and modulates downstream tumorigenesis regulators. H2009 and H1299 cells were treated with either scrambled control siRNA (SCR) or CyKILRb siRNA (siCyKILRb). Total RNA was extracted and subjected to deep RNA sequencing. After data filtering, clean-read alignment, gene expression analysis, and differential expression gene (DEG) analysis, KEGG enrichment analysis of annotated DEGs were performed using R Phyper. **A.** KEGG pathway enrichment analysis of differentially expressed genes following CyKILRb downregulation in H2009 and H1299 (Red arrow designates the PI_3_K-AKT signaling pathway with the largest enrichment). **B.** KEGG signaling pathway analysis of the PI_3_K-AKT comparing CyKILRb downregulation versus SCR in H2009 and H1299: upregulated genes = red box, downregulated genes = blue box. **C.** Heatmap depiction showing the top DEGs in the PI_3_K-AKT signaling pathway following CyKILRb downregulation versus SCR in H2009 and H1299 (red = upregulated genes; blue = downregulated genes). **D.** RNA expression levels of PIK3R2, RPS6KB2, GNB2, CDKN1A, and CDKN1B in H2009 and H1299 following CyKILRb downregulation compared to SCR. Data are presented as dot plots (n ≥ 3 replicates), and statistical analysis was performed using an unpaired t-test with Welch’s correction; ∗∗∗P < 0.001. **E.** Protein levels of PIK3R2, RPS6KB2, GNB2, CDKN1A (p21), and CDKN1B (p27) in H2009 and H1299 following CyKILRb downregulation compared to SCR analyzed using SDS-PAGE and western immunoblotting for the depicted factor. **F.** Quantification analysis of protein level normalized to GAPDH. Data are presented as dot plots (n = 3 replicates), and statistical analysis was performed using an unpaired t-test with Welch’s correction; ∗P < 0.05, ∗∗P < 0.01, ∗∗∗P < 0.001.

**Figure 2.**
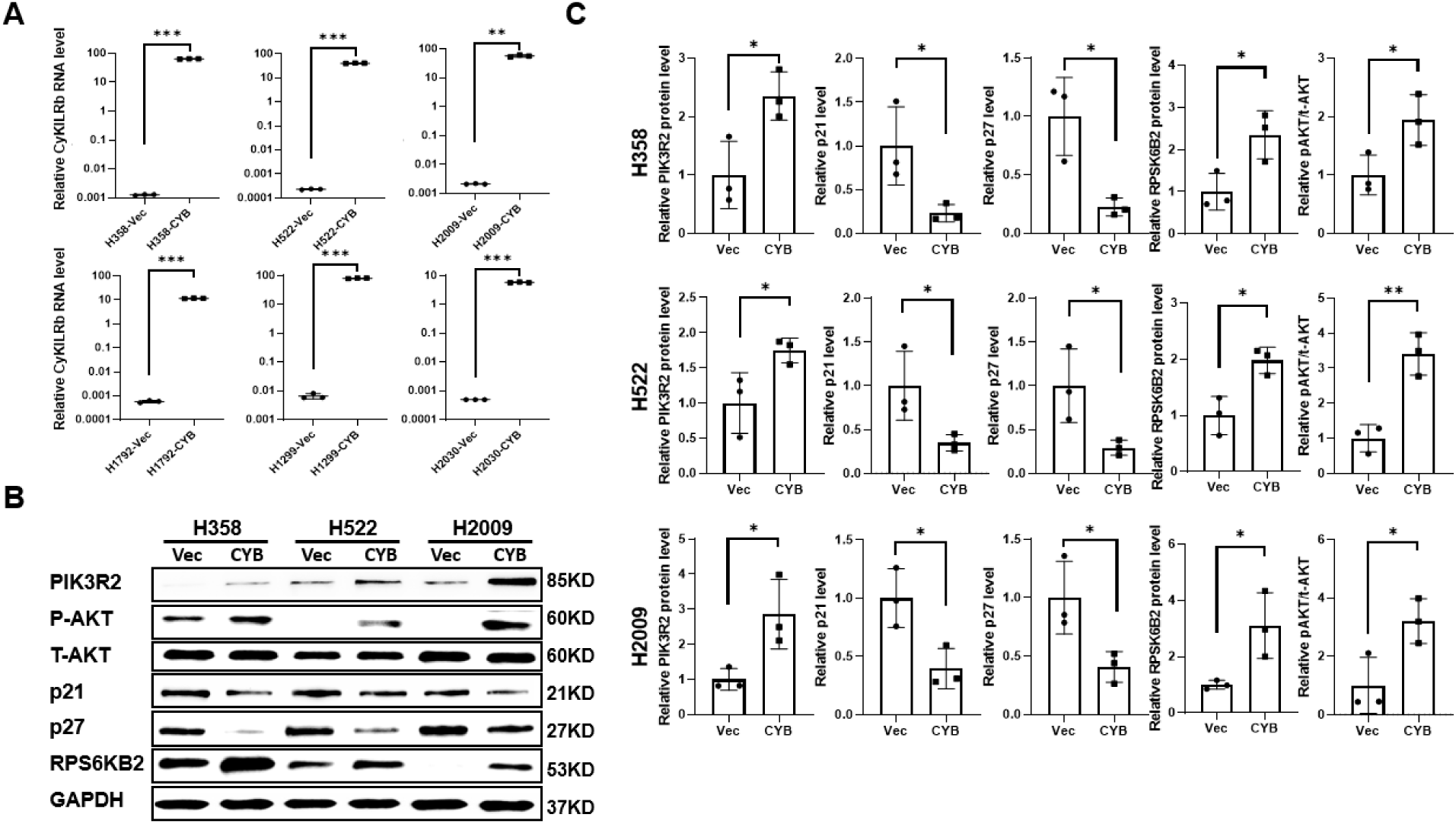
Ectopic expression of CyKILRb activates the PI_3_K-AKT pathway. The designated NSCLC cell lines were transfected with either an empty vector control (Vec) or CyKILRb (CYB) plasmids. Forty-eight hours post-transfection, the cells were subjected to: **A.** qRT-PCR analysis of CyKILRb RNA levels in the depicted six different NSCLC cell lines, and statistical analysis was performed using an unpaired t-test with Welch’s correction; ∗∗P < 0.01, ∗∗∗P < 0.001. **B.** SDS-PAGE and Western immunoblot analysis of key genes in the PI_3_K-AKT and tumorigenesis pathways for H358, H522, and H2009 cell lines ± empty vector control (Vec) or CyKILRb (CYB) plasmids. **C.** Quantification analysis of protein level normalized to GAPDH for the data depict in Figure B and repeated biological replicates. Data are presented as dot plots (n = 3 replicates), and statistical analysis was performed using an unpaired t-test with Welch’s correction; ∗P < 0.05, ∗∗P < 0.01.

### PIK3R2 downregulation mitigates CyKILRb augmentation of tumorigenic phenotypes

As CyKILRb induced AKT activation, we examined whether the PI_3_K activator, PIK3R2, whose expression was modulated by CyKILRb, was an upstream regulator of both AKT activation and downstream oncogenic phenotypes enhanced by this lncRNA. In contrast to our reported effects of CyKILRb downregulation in NSCLC cells [15], CyKILRb ectopic expression induced a significant increase in both cellular proliferation and clonogenicity (**Figure 3A&B**) as well as a significant increase in AKT phosphorylation and RPS6KB2 with concomitant loss of p21 and p27 (**Figure 3C**). Each of these parameters were significantly attenuated by PIK3R2 downregulation in comparison to controls (**Figure 3C-F**) with complete ablation of the enhanced clonogenicity imparted by CyKILRb expression (**Figure 3F**). Cellular proliferation was also suppressed by PIK3R2 downregulation as to fold enhancement by CyKILRb ectopic expression (∼50% on average across the three NSCLC cell lines examined), but the significant enhancement of this parameter by CyKILRb ectopic expression was still observed (**Figure 3D**). These data demonstrate that CyKILRb modulates the activation of AKT and downstream oncogenic phenotypes by regulating PI3KR2 expression.

**Figure 3.**
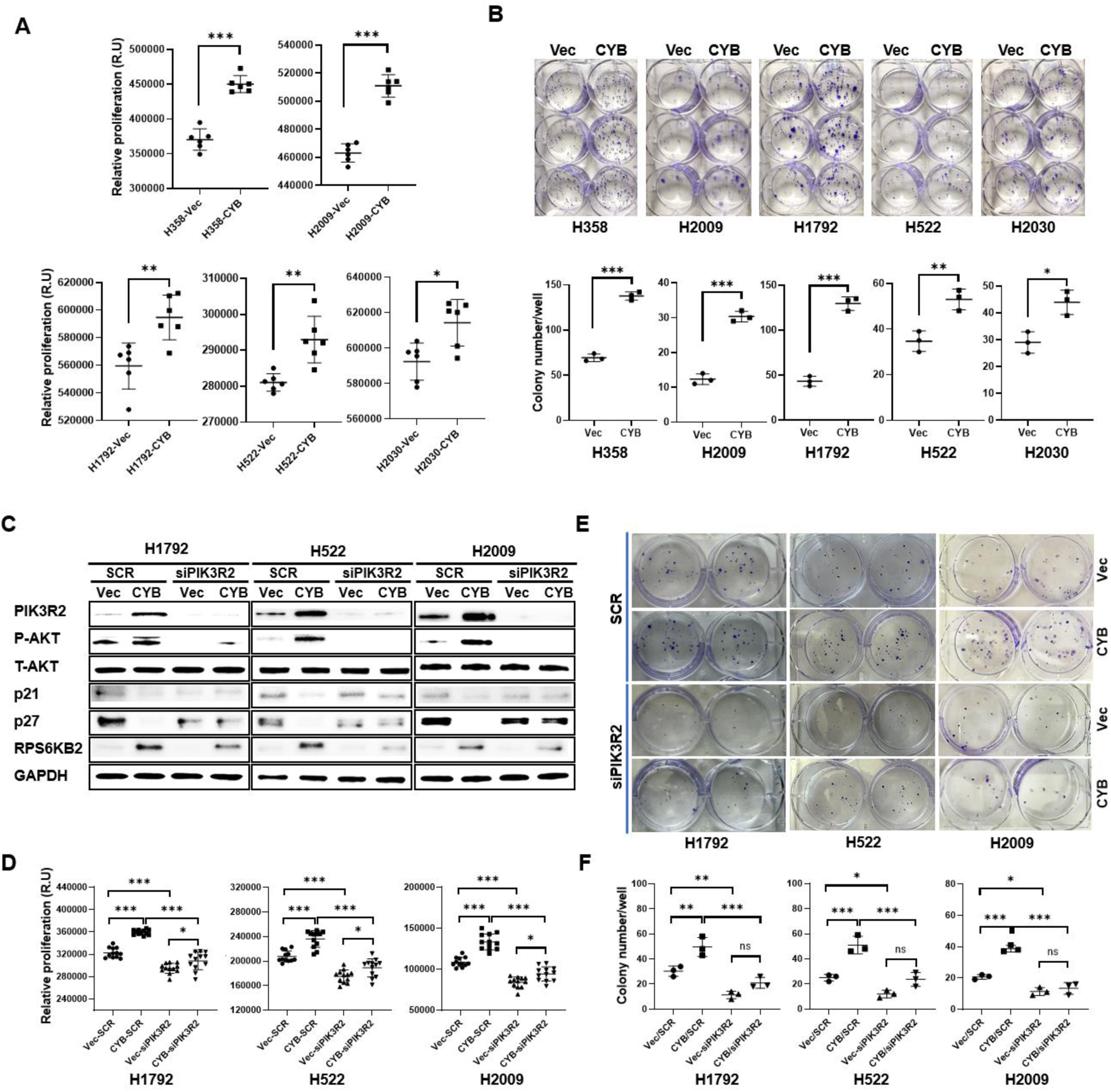
PIK3R2 gene downregulation mitigates CyKILRb effects on tumorigenic phenotypes. The depicted cell lines were transfected with either an empty vector control (Vec) or CyKILRb (CYB) plasmids. After 24 hours, cells were further transfected ± scrambled siRNA (SCR) or PIK3R2-specific siRNA (siPIK3R2). The depicted downstream analyses were conducted 24 hours post-siRNA transfection. **A.** Cell proliferation assay comparing CyKILRb ectopic expression (CYB) vs vector control (Vec). **B.** Clonogenic survival assay comparing CyKILRb ectopic expression (CYB) vs vector control (Vec). **C.** Protein levels of PIK3R2, phospho-AKT (P-AKT), total AKT (T-AKT), p21, p27, RPS6KB2, and GAPDH (for normalization) in H1792, H2009, and H522 cells (representative of 3 biological replicates). **D.** Proliferation analysis of Vec or CYB ± SCR or siPIK3R2 transfected samples in H1792, H2009, and H522 cells. **E.** Clonogenic survival analysis of Vec or CYB ± SCR or siPIK3R2 transfected samples in H1792, H2009, and H522 cells. **F.** Quantification of cell colonies in H1792, H2009, and H522 cells depicted in Panel E and replicates. For Panels A & B, data are presented as dot plots (n ≥ 3 replicates), and statistical analysis was performed using an unpaired t-test with Welch’s correction; ∗P < 0.05, ∗∗P < 0.01, ∗∗∗P < 0.001. For Panels D & F, data are presented as dot plots (n ≥ 3 replicates), and statistical analysis was performed using one-way ANOVA with Tukey’s multiple comparisons test; ns = not significant, ∗P < 0.05, ∗∗P < 0.01, ∗∗∗P < 0.001.

### CyKILRb enhances cell proliferation and clonogenic survival of NSCLC cells via the PI_3_K/AKT pathway

The above findings strongly suggested that CyKILRb enhanced NSCLC cell proliferation and clonogenicity via activation of the PI_3_K/AKT pathway as PIK3R2, a key regulator of PI_3_K activation, was downstream of CyKILRb in modulating these oncogenic phenotypes. To investigate this signaling link, PI_3_K or AKT were inhibited concurrently with CyKILRb ectopic expression. Inhibition of either the PI_3_K or AKT significantly suppressed the fold-induction of both NSCLC cell proliferation and clonogenic survival enhanced by CyKILRb ectopic expression (**Figure 4A-D**). Conversely, the ectopic expression of either constitutively active AKT (caAKT) (**Figure 5A-C**) or PIK3R2 (**Figure 6A-C**) “rescued” the suppression of both clonogenicity and proliferation caused by CyKILRb downregulation. Additionally, ectopic expression of PIK3R2 also “rescued” the loss of AKT phosphorylation caused by CyKILRb downregulation (**Figures 6A**). These data demonstrate that CyKILRb drives cell proliferation and clonogenicity via the PI_3_K/AKT axis and further orient the signaling pathway from CyKILRb → PIK3R2 → PI_3_K/AKT.

**Figure 4.**
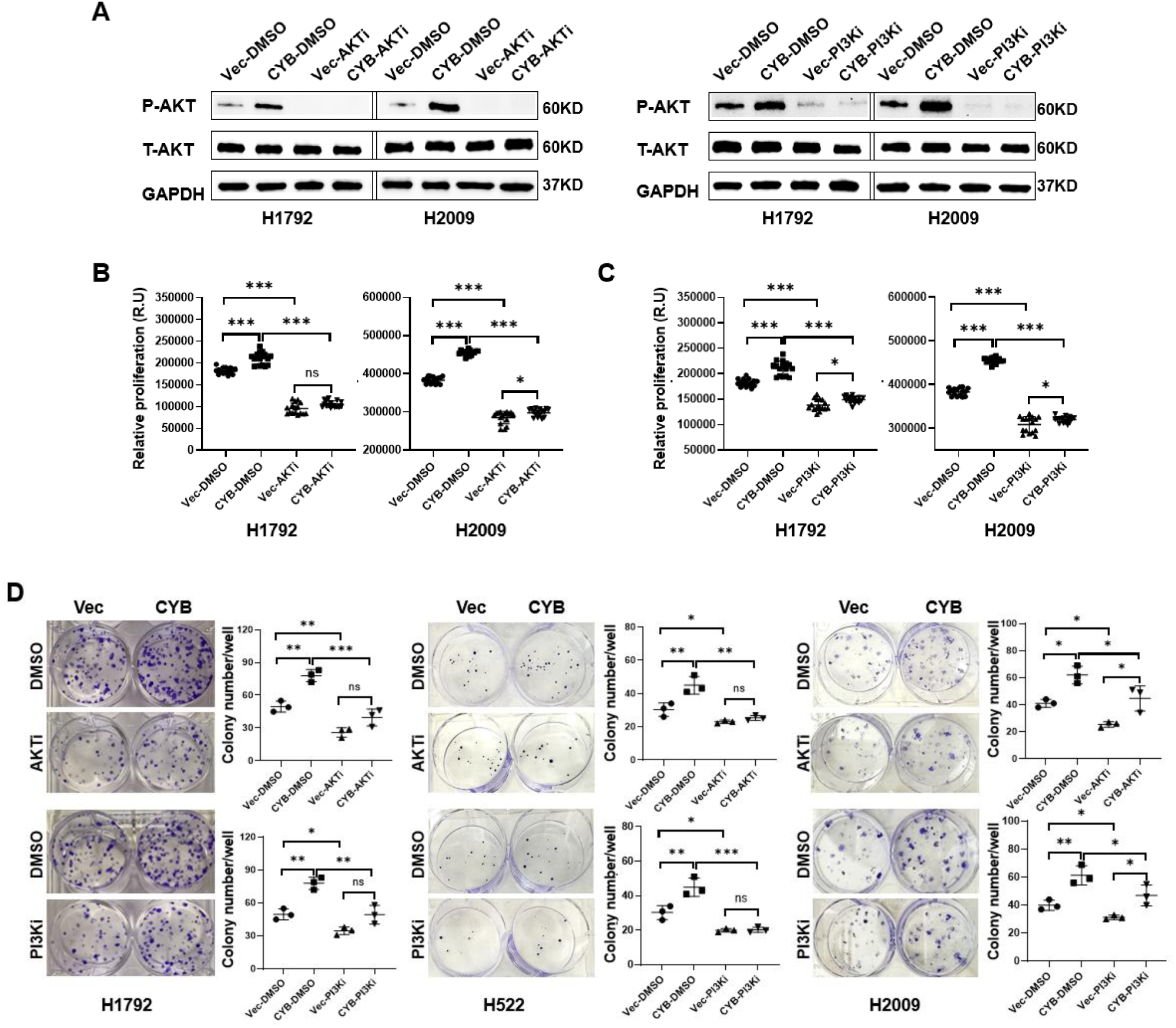
The PI_3_K/AKT pathway is a key downstream mediator of CyKILRb-induced tumorigenic phenotypes. The depicted cell lines were transfected with either an empty vector control (Vec) or CyKILRb (CYB). After 24 hours, cells were further treated ± a PI3K (PI3Ki) or AKT inhibitor (AKTi). After an additional 24 hours, the following analyses were performed: **A.** SDS-PAGE and Western immunoblotting for phospho-AKT (P-AKT), total AKT (T-AKT), and GAPDH (for normalization) of Vec or CYB ± AKT inhibitor or PI_3_K inhibitor treatment in H1792 and H2009 cells (representative of 3 biological replicates). **B.** Proliferation analysis of Vec or CYB ± AKT inhibitor-treatment in H1792 and H2009 cells. **C.** Proliferation analysis of Vec or CYB ± PI_3_K inhibitor treatment in H1792 and H2009 cells. **D.** Clonogenic survival analysis and quantification of Vec or CYB ± AKT inhibitor or PI_3_K inhibitor treatment in H1792, H522, and H2009 cells. Data depicted in Panels B-D are presented as dot plots (n ≥ 3 replicates), and statistical analysis was performed using one-way ANOVA with Tukey’s multiple comparisons test (n ≥ 3 replicates); ns = not significant, ∗P < 0.05, ∗∗P < 0.01, ∗∗∗P < 0.001.

**Figure 5.**
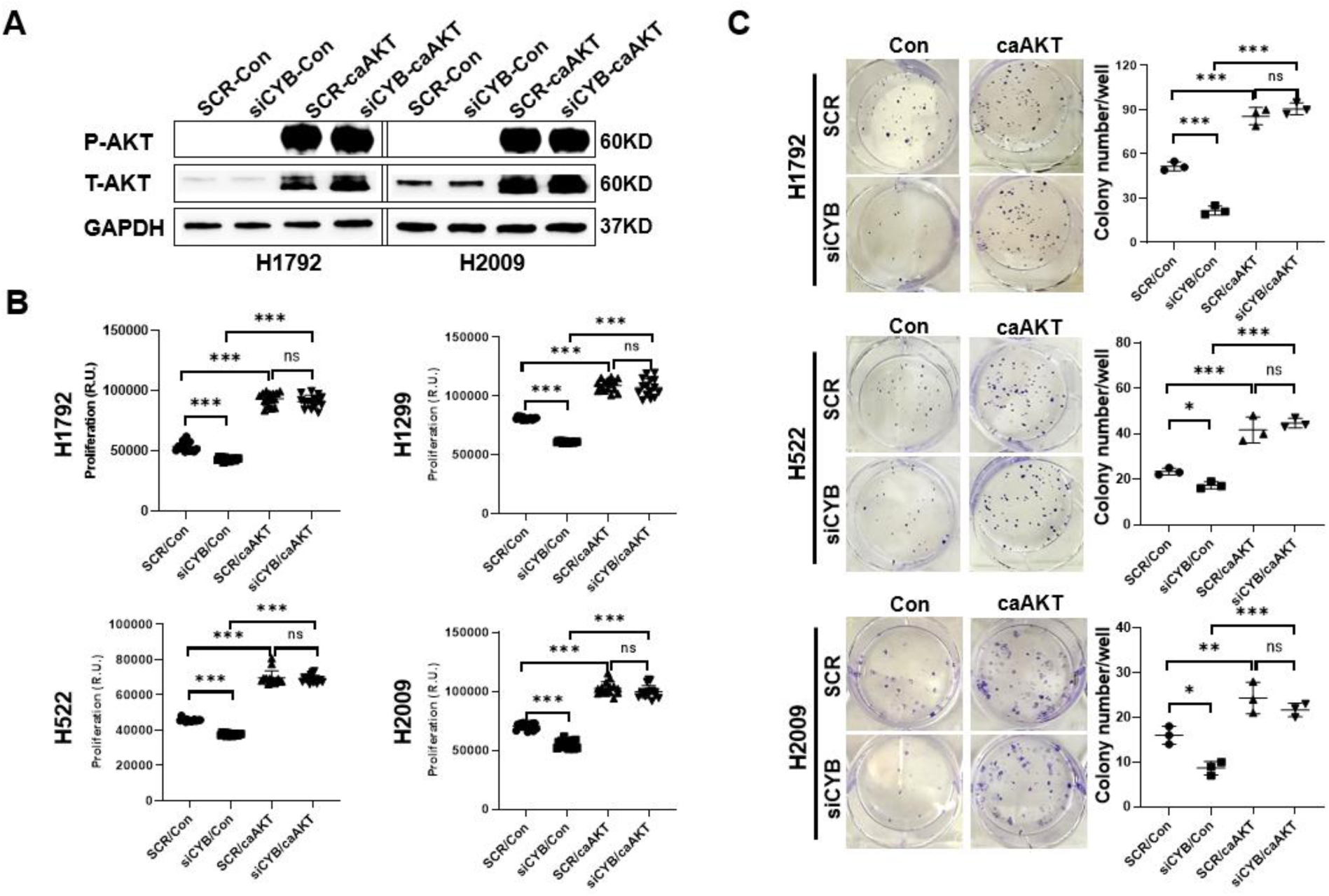
Ectopic expression of constitutively-active AKT rescues the suppression of tumorigenic phenotypes caused by CyKILRb loss. Cells were transfected with either scrambled siRNA (SCR) or CyKILRb-specific siRNA (si-CYB). After 24 hours, cells were transfected with an empty vector (Vec) or constitutive-active AKT (ca-AKT) expression constructs. The following downstream analyses were then performed 24 hours post-plasmid transfection: **A.** SDS-PAGE and Western immunoblotting for the protein levels of P-AKT, T-AKT, and GAPDH in H2009 and H1792 cells (representative of 3 biological replicates). **B.** Cell proliferation analysis in H522, H2009, H1299, and H1792 cells transfected with SCR or si-CYB ± Vec or caAKT. **C.** Clonogenic survival assay in H1792 and H522 cells transfected with SCR or si-CYB ± Vec or caAKT. Data depicted in Panels B and C are presented as dot plots (n ≥ 3 replicates), and statistical analysis was performed using one-way ANOVA with Tukey’s multiple comparisons test (n ≥ 3 replicates); ns = not significant, ∗P < 0.05, ∗∗P < 0.01, ∗∗∗P < 0.001.

**Figure 6.**
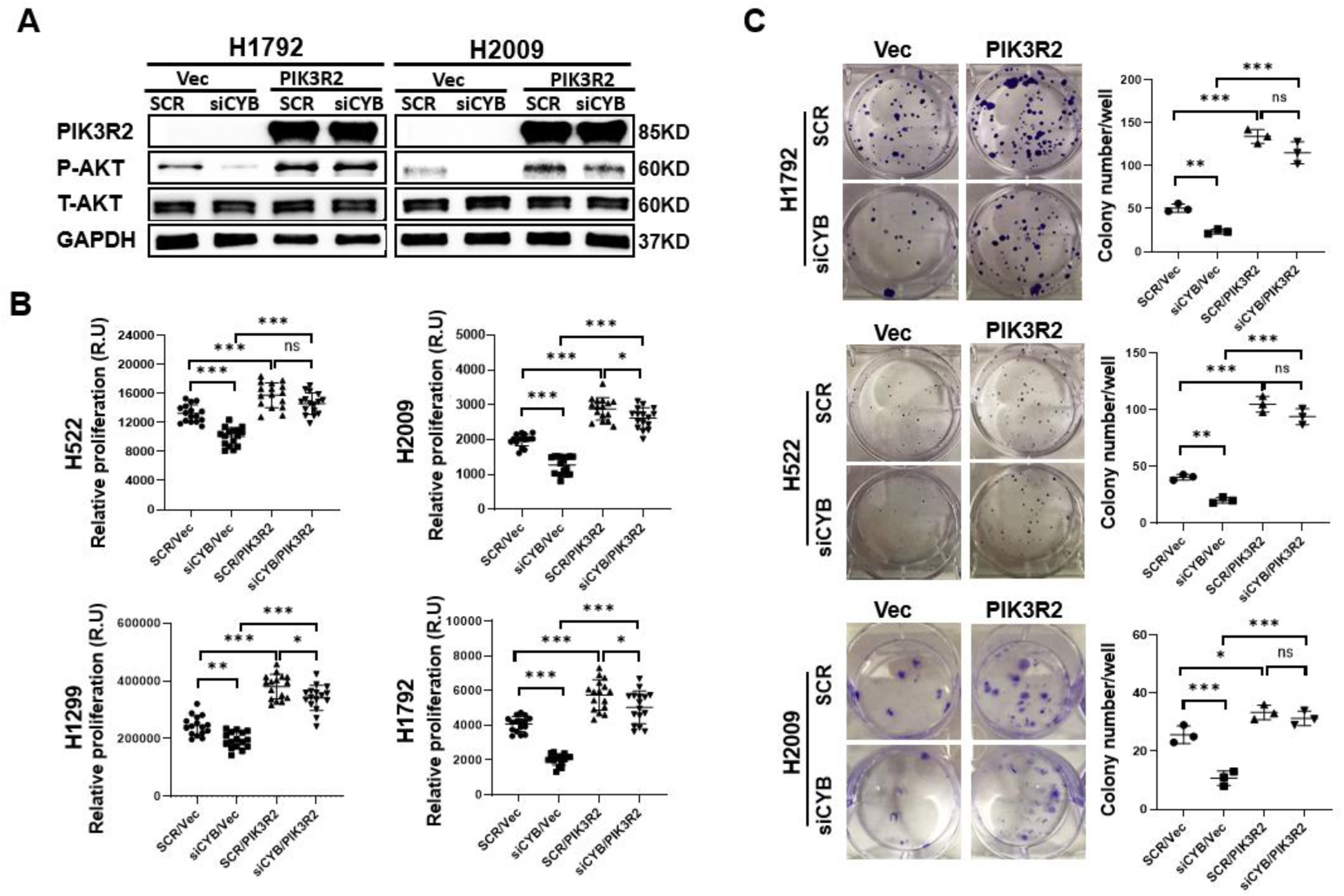
PIK3R2 is a downstream mediator of CyKILRb in driving the PI_3_K/AKT pathway and cell survival/clonogenicity. Cells were transfected with either scrambled siRNA (SCR) or CyKILRb-specific siRNA (siCYB). After 24 hours, cells were transfected with an empty vector (Vec) or PIK3R2 expression constructs. The following downstream analyses were then performed 24 hours post-plasmid transfection **A.** SDS-PAGE and Western immunoblotting for the protein levels of P-AKT, T-AKT, PIK3R2, and GAPDH in H2009 and H1792 cells (representative of 3 biological replicates). **B.** Cell proliferation analysis in H522, H2009, H1299, and H1792 cells transfected with SCR or siCYB ± Vec or PIK3R2. **C.** Clonogenic survival assay in H1792, H2009, and H522 cells transfected with SCR or siCYB ± Vec or PIK3R2. **D.** Quantification of colonies formed in the clonogenic survival assays depicted in Panel C and replicates. Data depicted in Panels B and C are presented as dot plots (n ≥ 3 replicates), and statistical analysis was performed using one-way ANOVA with Tukey’s multiple comparisons test (n ≥ 3 replicates); ns = not significant, ∗P < 0.05, ∗∗P < 0.01, ∗∗∗P < 0.001.

### CyKILRb activation of the PI_3_K/AKT axis negatively regulates the inclusion of CyKILR exon 3 blocking the expression of the tumor suppressor lncRNA, CyKILRa

The nuclear CyKILR splice variant, CyKILRa, in stark contrast to CyKILRb, is a strong tumor suppressor, and our laboratory previously reported that CyKILRb suppressed the expression of CyKILRa[15]. To determine whether a reciprocal regulatory relationship exists between the two CyKILR splicing variants via signaling pathways downstream of CyKILRb, we analyzed the expression of both CyKILR splice variants upon ectopic expression of CyKILRb. The ectopic expression of CyKILRb induced a significant reduction in the inclusion of exon 3 into the mature CyKILR mRNA leading to a loss in CyKILRa and sustained expression of CyKILRb (**Figure 7A&B**). Since CyKILRb modulates the PI_3_K/AKT pathway, we investigated whether inhibition of PI_3_K or AKT influences CyKILRa levels. Our results showed that CyKILRa levels increased upon inhibition of either PI_3_K or AKT (**Figure 7C-F**). Notably, ectopic PIK3R2 expression also abolished the upregulation of CyKILRa induced by CyKILRb downregulation (**Figure 7G&H**). Similar to PIK3R3 downregulation, PI_3_K or AKT inhibition augmented the expression of p27 and p21 (**Figure 7C&E**). Taken together, these findings demonstrate that CyKILRb induces PIK3R2, thereby activating the PI_3_K/AKT pathway to suppress known the tumor suppressors, CyKILRa, p27, and p21, which correlates with enhanced cell proliferation and clonogenic potential.

**Figure 7.**
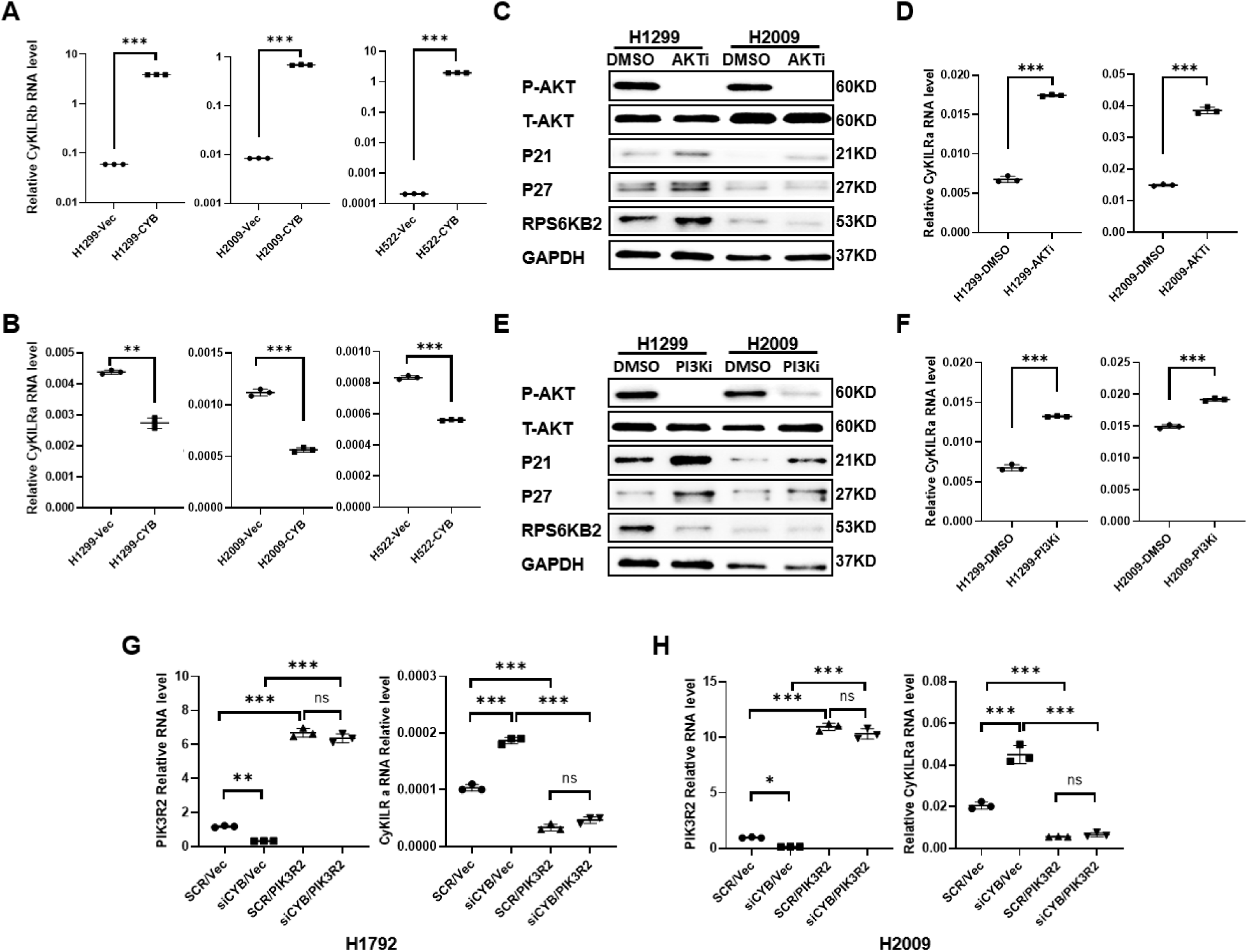
CyKILRb negatively regulates the inclusion of CyKILR exon 3 suppressing CyKILRa expression though the PI_3_K-AKT pathway. **A, B.** The depicted cell lines were transfected with either an empty vector control (Vec) or CyKILRb (CYB) plasmids. 24 hours post-transfection, CyKYLRb (**A**) and CyKILRa (**B**) transcript levels were determined via qRT-PCR. **C-F.** The depicted cell lines were treated ± a PI3K inhibitor (PI3Ki) or AKT inhibitor (AKTi). After 24 hours, the following downstream analyses were then performed: **C,E.** SDS-PAGE and Western immunoblotting for the protein levels of P-AKT, T-AKT, p21, p27, RPS6KB2, and GAPDH. **D,F.** CyKILRa transcript levels were determined via qRT-PCR. Data depicted in Panels A, B, D, and F are presented as dot plots (n ≥ 3 replicates), and statistical analysis was performed using an unpaired t-test with Welch’s correction; ∗P < 0.05, ∗∗P < 0.01, ∗∗∗P < 0.001. **G,H.** Cells were transfected with either scrambled siRNA (SCR) or CyKILRb-specific siRNA (siCYB). After 24 hours, cells were transfected with an empty vector (Vec) or PIK3R2 expression constructs. 24 hours post-plasmid transfection,PIK3R2 and CyKILRa transcript levels were determined via qRT-PCR. Data are presented as dot plots (n ≥ 3 replicates), and statistical analysis was performed using one-way ANOVA with Tukey’s multiple comparisons test (n ≥ 3 replicates); ns = not significant, ∗P < 0.05, ∗∗P < 0.01, ∗∗∗P < 0.001.

## MATERIALS AND METHODS

### Cell lines and materials

NCI-358, NCI-H522, NCI-H2009, NCI-H2030, NCI-H1299, and NCI-H1792 were purchased from ATCC. SuperScript™ IV VILO™ Master Mix with ezDNase™ Enzyme (11766050), TaqMan™ Universal PCR Master Mix (4324018), Opti-MEM™ I Reduced Serum Medium (31985062), Lipofectamine™ RNAiMAX Transfection Reagent (13778150), Lipofectamine™ 3000 Transfection Reagent (L3000015), and Pierce Fast Western Blot Kit (35050) were from Thermo Fisher Scientific. Cell Titer-Glo 2.0 Assay Kit was purchased from Promega (G9242). The mycoplasma contamination detection kit was purchased from InvivoGen (rep-mys-50).

### Cell culture

All cells were cultured following the procedures previously described and were utilized within 6 months or -twenty passages, whichever came first, from the time of receipt and thawing[15]. Throughout the entire study, cell lines were subjected to Mycoplasma testing every two months using a Mycoplasma contamination detection kit. Parental cell lines were authenticated by STR (short tandem repeat) profiling.

### RNA extraction and Quantitative RT-PCR

Total RNA was isolated from cultured cells using the RNA Subcellular Isolation Kit from Active Motif (25501) following the manufacturer’s instructions. RNA quantification was performed using a NanoDrop NP-1000 spectrometer, and RNA (2 μg) was used for the first-strand cDNA synthesis using SuperScript™ IV VILO™ Master Mix with ezDNase™ Enzyme. Subsequently, qRT-PCR was conducted as done on a Quant Studio 5 instrument (Applied Biosystems, Carlsbad, CA, USA) [16–23]. The housekeeping gene GAPDH was utilized for normalization using the ΔΔCt method. The detection of CyKILRa follows the methods a reported by us [15].

### siRNA and plasmid transfection

Cells were plated in a full medium without antibiotics 24hrs before transfection. Cells were transfected at 60% confluency. siRNA or plasmid DNA were diluted in Opti-MEM® I Reduced Serum Medium. For siRNA transfection, Lipofectamine RNAiMAX was employed as done by us[15] , while Lipofectamine 3000 was used for plasmid transfection.[24–28] The final concentration of siRNA was 20 nM in the cell culture medium. For CyKILRb transfection, 3 μg plasmid and 6 ul lipofectamine 3000 were used for 1 million cells in 6 ml of cell culture medium. For PIK3R2 and ca-AKT transfection, 12 μg plasmid and 30 μl lipofectamine 3000 were used for 1-1.5 x10^6^ cells in 6 ml cell culture medium[29].

### PI_3_K and AKT inhibitor treatment

Cells were cultured in full medium until reaching 60% to 80% confluency before inhibitor treatment. Cells were treated with PI_3_K inhibitor (LY294002, Tocris) at 6 μM or AKT inhibitor (Akt Inhibitor VIII, Cayman) at 10 μM for 6 hours for protein assay and 24 hours for RNA analysis. Subsequently, the full medium was replaced with serum-free medium when treated with inhibitor. After treatment, full medium was replaced and culture for an additional 24-36 hours before for proliferation assay.

### Proliferation analysis

To investigate cell proliferation, the Cell Titer-Glo Luminescent Cell Viability kit was utilized following the manufacturer’s instructions. Cells were initially seeded in 96-well plates at a density of 2000 cells per well in RPMI1640 supplemented with 10% FBS. After treatment with CyKILRb, PIK3R2, ca-AKT or siRNA or inhibitor for 48 hours, the plates were carefully decanted and refilled with 50 μl of Cell Titer-Glo reagent and 50 μl of PBS per well. The plates were then incubated for 10 minutes at room temperature in the dark. Subsequently, the luminescence readings were taken using the Gene5 Multi-Mode Microplate Reader (Molecular Devices, Sunnyvale, CA) with a 250 ms integration.

### Clonogenic survival assays

After plasmid transfection or siRNA treatment, cells were harvested and re-suspended in complete growth media. Subsequently, the cells (200-400) were seeded onto six-well plates and allowed to grow until visible colonies formed as done, which typically occurred within 7-21 days depending on cell type.[16–19, 21, 22, 30, 31]. Once the colonies became visible, they were fixed with cold 20% methanol, stained with 0.2% crystal violet in 20% methanol, washed, and then left to air-dry as previously done by us[15].

### Western immunoblotting

Cells were lysed in RIPA buffer containing Halt™ Protease and Phosphatase Inhibitor Cocktail (Thermo Fisher Scientific, 78440). For protein concentration determination, the Pierce BCA Protein Assay kit (Thermo Fisher Scientific, 23227) was used.[20, 32–35] Whole-cell lysates were mixed with Laemmli loading buffer, boiled, separated by 4-20% SDS–PAGE, and transferred to a PVDF membrane as done[32, 34, 36–39]. Subsequently, immunoblot analyses were performed using the following primary antibodies: anti-PIK3R2/p85 (cat. no. 83606-5-RR; 1:2000), p21 (cat. no. 10355-1-AP; 1:1000), p27 (cat no. 25614-1-AP; 1:1000), RPS6KB2 (cat. no. 26194-1-AP; 1:500), GNB2 (cat. no. 16090-1-AP; 1:500), and GAPDH (cat. no. 60004-1-Ig; 1:50,000) from ProteinTech; anti-PI3KB/p110 (cat. no. ab151549; 1:1000) from Abcam; and anti-phospho-AKT (Ser473; cat. no. 4060S; 1:1000) and AKT (pan; Cell cat. no. 8596S; 1:2000) from Signaling Technology. The specific protein bands were visualized by using Western Blotting Kit (Pierce, 35050).

### Transcriptomics and Signaling Pathway Analyses

RNA sequencing was conducted using 150 bp paired-end reads, aiming for a depth of approximately 30 million reads as reported by us[15] . The sequencing data were filtered with SOAPnuke (v1.5.6) (removing reads containing sequencing adapter and low-quality base ratio), then the clean reads were stored in FASTQ format.[40] Differential expression gene (DEG) analysis was performed using the DESeq2 with Q value ≤ 0.05.[41] KEGG enrichment analysis of annotated different expression gene was performed by R Phyper (v120.0) based on Hypergeometric test. The significant levels of terms and pathways were corrected by Q value with a rigorous threshold (Q value ≤ 0.05).

### Statistical analysis

Graphing and statistics were performed using Prism GraphPad (Prism Software, San Diego, CA). The following models were used for statistical analysis as described, unpaired t-test with Welch’s correction were used when only two experimental groups were being compared[42]. Repeated measures ANOVA were used when analyzing comparison groups that contained a dependent variable that was measured several times over the course of the experiment[43]. One-way ANOVA with Tukey’s multiple comparisons test were used when comparing three or more independent groups[44, 45]. All data reported as dotplot or mean (SD) where applicable; p < 0.05 was considered statistically significant. All bioinformatic analyses were conducted using the R program (NS = not significant, ∗P < 0.05, ∗∗P < 0.01, ∗∗∗P < 0.001).

## DISCUSSION

Using an unbiased molecular approach, this study uncovered a novel oncogenic mechanism in non-small cell lung cancer (NSCLC) driven by the cytoplasmic long non-coding RNA splice variant, CyKILRb. In particular, this study demonstrated that CyKILRb promotes clonogenicity and cellular proliferation by activating the PI_3_K/AKT signaling pathway, primarily through upregulation of phosphoinositide-3-kinase regulatory subunit 2 (PIK3R2). PIK3R2 encodes the regulatory subunit (p85β) of PI_3_K, a well-established mediator of signaling pathways crucial for cell growth and survival[46]. Mutations in PIK3R2 have been linked to various developmental brain disorders, including megalencephaly-polymicrogyria-polydactyly-hydrocephalus (MPPH) syndrome and bilateral perisylvian polymicrogyria (BPP)[47]. In cancer, the role of PIK3R2 is understudied, but the factor is considered oncogenic, particularly in certain cancers such as glioblastoma multiforme[48]. Our study supports a role for PIK3R2 as an oncogenic factor, and in the context of NSCLC and CyKILRb expression, a major downstream driver of the PI_3_K/AKT pathway, which fosters key cancer cell phenotypes. Thus, PIK3R2 expression is a plausible biomarker for constitutively activation of the PI_3_K/AKT pathway in the absence of a PI_3_K mutation. As this pathway is hyperactive in 50-73% of NSCLC with major consequences on oncogenesis, the targeting of PIK3R2 and CYKLRb expression becomes a plausible therapeutic option[49–51]. Additionally, the high expression of either factor may guide precision-based therapies for use of AKT inhibitors, which has been a large hurdle for determining the patient population that may benefit from these therapies[52].

Although understudied, recent reports have shown that the expression of PIK3R2 is regulated by microRNAs (miRs). More specifically, microRNA-126 (miR-126) targets PIK3R2 and suppresses the proliferation of NSCLC cells[53]. Additionally, PIK3R2 expression has been linked to loss of miR-30a-5p and constitutively active EGFR signaling as well as resistance to tyrosine kinase inhibitors (TKIs) in NSCLC[54]. More recently, co-targeting of miR-126-3p and miR-221-3p was shown to inhibit PIK3R2 reducing lung cancer growth and metastasis. These studies suggest that PIK3R2, in particular, is highly regulated by miRs in NSCLC, and our laboratory recently reported that CyKILRb may act as a direct “sink/sponge” or indirect regulator of several important miRs for the growth and clonogenicity of NSCLC cells such as miR-424–3P and miR-503-3P[15]. Due to the strong link between CyKILRb and PIK3R2 expression, it is highly plausible that PIK3R2 levels are enhanced by CyKILRb through modulation of miRs. Indeed, the lncRNA, HOTAIR, has been reported to modulate PIK3R2 levels via acting as a “sponge” for additional PIK3R2 targeting miRs[55]. Future studies to delineate specific CyKILRb/miR interactions are thus warranted and have potential clinical applications.

In this study, downregulation of CyKILRb led to the decreased expression of PIK3R2 and downstream signaling targets, including phosphorylated AKT, while increasing the expression of tumor suppressors, p21 and p27. These changes were associated with reduced cell proliferation and clonogenic survival. The link to p21 and p27 is highly relevant as these factors are well established regulators of NSCLC cell proliferation[56–60]. Our data support the activation of the PI_3_K/AKT pathway by CyKILRb as an upstream regulator of the expression of these tumor suppressors, which is in line with reports on the AKT pathway restraining the function of these two tumor suppressors[61]. For example, the AKT pathway regulates p27 expression in NSCLC by phosphorylating and inactivating the FOXO transcription factor, which normally promotes p27 transcription. AKT can also directly phosphorylate p27, which promotes its cytoplasmic localization. This ability of hyperactivated AKT signaling in NSCLC to reduce nuclear p27 contributes to increased cell proliferation, growth, and malignant progression. In regard to p21, hyperactivated AKT signaling in NSCLC has also been shown to lead to the downregulation of p21 contributing to increased cell proliferation in some instances[59, 62]. Similar to p27, AKT signaling can also influence p21 function through post-translational modifications. For example, phosphorylation of p21 by AKT induces cytoplasmic localization analogous to p27, which functionally inactivates it and promotes a proliferative effect[56]. Thus, CyKILRb may be a key upstream and previously overlooked regulator of these pathways.

Mechanistically, we also identified a novel mechanism in which CyKILRb downregulates the expression of the nuclear splice variant, CyKILRa, a known tumor suppressor, through PI_3_K/AKT signaling. Inhibition of either PI_3_K or AKT led to CyKILRa re-expression, establishing a signaling axis whereby CyKILRb promotes tumorigenesis by both activating growth-promoting pathways and silencing tumor suppressive isoforms via modulation of the alternative mRNA processing of the CyKILR pre-mRNA. This occurs via a novel feed-forward mechanism requiring suppression of CyKILR exon 3 inclusion, which sustains CyKILRb expression at the expense of CyKILRa. Previously, our laboratory demonstrated that the PI_3_K axis regulated the pre-mRNA processing of survival and apoptotic factors (BCL-X and caspase 9) in NSCLC cells [15, 19, 20, 30, 63–65]. Whereas BCL-X RNA splicing was linked to PKCι downstream of PI_3_K, caspase 9 RNA splicing was regulated by AKT downstream of PI_3_K, which is a similar model to our findings in this study. In the specific case of caspase 9 RNA splicing, AKT directly phosphorylated hnRNP L, which suppressed the inclusion of an exon cassette. Whereas how CyKILR RNA splicing is regulated has not been determined, the suppression of CyKILR exon 3 inclusion via AKT is highly suggestive of a similar mechanism to caspase 9 RNA splicing.

Also of note in this study is that ectopic expression of PIK3R2 did not fully restore the tumorigenic phenotypes lost when CyKILRb was downregulated. These data suggest that while PIK3R2 is a key downstream effector of CyKILRb, it is not the sole mediator. CyKILRb may regulate additional targets or interact with RNA-binding proteins, mRNA stability factors, or other signaling molecules that contribute independently to oncogenesis via separate signaling pathways. Moreover, the unique RNA structure and cytoplasmic localization of CyKILRb could facilitate scaffolding or decoy functions that extend beyond PIK3R2 activation and are not recapitulated by PIK3R2 overexpression alone. In this regard, our pathway analysis of deep RNA sequencing data did show strong effects on other known oncogenic pathways such as the JAK-STAT pathway[66]. Thus, CyKILRb may act as a broad regulator of multiple survival and oncogenic pathways making this lncRNA an intriguing anti-cancer target in the context of specific oncogenotypes in NSCLC. Additionally, these findings suggest that the exclusive expression of CyKILRb in NSCLC with wild-type and active STK11 and CDKN2A is required to overcome these tumor suppressive pathways and maintain a tumorigenic phenotype.

In summary, our findings define CyKILRb as a critical modulator of the PI_3_K/AKT signaling cascade and as a repressor of its nuclear tumor-suppressive counterpart, CyKILRa. This dual function positions CyKILRb as a potent oncogenic driver in NSCLC. The partial rescue by PIK3R2 overexpression emphasizes the multifaceted nature of lncRNA-mediated signaling, where alternative splicing, cellular localization, and RNA structure contribute to diverse functional outputs. Targeting CyKILRb or its broader network may offer novel therapeutic strategies for NSCLC patients, especially those tumors exhibiting hyperactivation of the PI_3_K/AKT pathway with wild-type CDKN2A and STK11 genes.

## Data and Materials Availability

All RNAseq data are in the process of submission to NCBI Sequence Read Archive (SRA) database for public access (Submission number: SUB15688856).

## Acknowledgements

The contents of this manuscript do not represent the views of the Department of Veterans Affairs or the United States Government. This work was mainly supported by the Veteran’s Administration (VA Merit Review, BX001792 (CEC), VA Merit Review award, BX 006063 (CEC), VA Merit Review award, RD001334 (CEC), and a Senior Research Career Scientist Award, IK6BX004603 (to CEC)). This work was peripherally supported by way of research cores, methodology and technology development, and software development by the National Institutes of Health by way of P01 CA171983 (CEC) and P01 CA302570 (CEC), NIH/NCI Cancer Center Support Grant P30 CA044579, and the R01s AI139072 (to CEC), and DK126444 (to CEC). This work was also supported by funds from the Department of Medicine in the University of Virginia (UVA)-School of Medicine as well as the UVA NCI Comprehensive Cancer Center in regard to software licensing and equipment purchases.

## Author Contributions

XX – designing research studies, conducting experiments, acquiring data, analyzing data, and writing the manuscript. HPM – developing and analyzing short and long read deep RNA sequencing data, manuscript method writing, and figure composition. ALL – conducting experiments, acquiring data, and analyzing data. CEC – overall supervision, designing research studies, analyzing data, providing reagents and funding, editing manuscript, and writing the manuscript.

